# Computationally replicating the Smith et al. (2015) positive-negative mode linking functional connectivity and subject measures

**DOI:** 10.1101/2020.04.23.058313

**Authors:** Nikhil Goyal, Dustin Moraczewski, Peter Bandettini, Emily S. Finn, Adam Thomas

**Affiliations:** Data Science and Sharing Team, National Institute of Mental Health, Bethesda, MD; Section on Functional Imaging Methods, National Institute of Mental Health, Bethesda, MD

## Abstract

Understanding brain functionality and predicting human behavior based on functional brain activity is a major goal of neuroscience. Numerous studies have been conducted to investigate the relationship between functional brain activity and attention, subject characteristics, autism, psychiatric disorders, and more. By modeling brain activity data as networks, researchers can leverage the mathematical tools of graph and network theory to probe these relationships. In their landmark study, Smith et al. (2015) analyzed the relationship of young adult connectomes and subject measures, using data from the Human Connectome Project (HCP). Using canonical correlation analysis (CCA), Smith et al. found that there was a single prominent CCA mode which explained a statistically significant percentage of the observed variance in connectomes and subject measures. They also found a strong positive correlation of 0.87 between the primary CCA mode connectome and subject measure weights. In this study, we computationally replicate the findings of the original study in both the HCP 500 and HCP 1200 subject releases. The exact computational replication in the HCP 500 dataset was a success, validating our analysis pipeline for extension studies. The extended replication in the larger HCP 1200 dataset was partially successful and demonstrated a dominant primary mode.

## 1. Introduction

Understanding the relationship between behavior, cognition, and functional brain organization is a major goal of neuroscience (S. Smith, 2016). Consequently, neuroscience research has become increasingly focused on investigating the relationship between whole-brain functional connectivity (i.e., the “connectome”) and human behavior (S. Smith, 2016; Sporns et al., 2005). A number of successful studies (Bassett & Sporns, 2017) have investigated the link between brain network activity and attention (Rosenberg et al., 2016), subject characteristics (S. M. Smith et al., 2015b), autism (Uddin, 2015), psychiatric disorders (Lydon-Staley & Bassett, 2018; Mier & Kirsch, 2015), and gene expression (Miller et al., 2016).

A notable study in this field (S. M. Smith et al., 2015b) investigated the relationship between functional connectivity and behavioral and phenotypic measures, using data from the Human Connectome Project (HCP) (Van Essen et al., 2012). Smith et al.’s 2015 analysis of data from the HCP 500 subject release (https://www.humanconnectome.org/study/hcp-young-adult/document/500-subjects-data-release) found a correlation between subject measures (SMs), such as lifestyle, demographics, and psychometric characteristics, and patterns of functional connectivity between brain regions.

Using canonical correlation analysis (CCA), the authors discovered one dominant CCA mode that explained substantially more of the observed covariance between functional connectivity and subject measures (SMs) than any other CCA mode. From this single mode, they derived a ‘positive-negative’ axis which quantified and depicted the correlation of various SMs to the CCA mode, linking lifestyle, demographic, and psychometric subject measures to each other and a specific pattern of brain connectivity (S. M. Smith et al., 2015b). SMs commonly considered positive qualities (e.g. higher income, greater education level, high performance on cognitive tests) were positively correlated with the primary mode, and SMs commonly considered negative qualities (e.g. substance use and anger) were negatively correlated with the primary mode. The primary mode of the CCA explained 0.53% of the variance in connectome edge weights, and approximately 1.7% of the variance in the subject measures, significantly outperforming all other CCA modes calculated. The primary mode’s z-scores were 7.7 and 9.2 for connectomes and SMs, respectively. The next highest z-scores (for any of the 99 other modes) were 2.7 and 2.4.

In addition to the positive-negative axis, Smith et al. analyzed specific region-to-region connections that varied as a function of the primary CCA mode, finding an overall positive correlation of *r=0.2* between the CCA connectome weights and the original group average functional connectome. A hierarchical clustering analysis of the 200 connectome nodes revealed clusters of nodes in four regions: one cluster in the sensory, motor, insula, and dorsal attention regions; and three clusters in covering the default mode network, subcortical, and cerebellar regions (S. M. Smith et al., 2015b). The brain areas that contributed strongest to variations in connectivity were similar to those associated with the default mode network (DMN), suggesting that functional connectivity within (and, to some degree, between) the DMN may be important for higher-level cognition and behavior (Wang et al., 2018).

Given the importance of the Smith et al. study and growing efforts toward reproducible science (Munafò et al., 2017), this preprint presents an exact computational replication of Smith et al.’s findings using data from the Human Connectome Project 500 subject release. This aspect of our study served two purposes: as an independent replication of a landmark finding in neuroscience, and to validate our replication pipeline for the purpose of extending this analysis to independent datasets outside the HCP study.

In addition, we performed a separate analysis in which we use data from the HCP 1200 subject release in the CCA analysis to determine if the single prominent mode phenomenon replicates in a larger sample of subjects from the same study. Because the HCP 1200 dataset is similarly homogenous as the HCP 500 dataset, we hypothesized that the single prominent mode would replicate in the HCP 1200 dataset. This aspect of our study was purely explorational and was not used to validate our pipeline.

The definition of a successful replication in the HCP 500 analysis was:

1. At least one CCA mode that explains a statistically significant amount of variance in the connectomes and subject measures, relative to the null distribution generated via permutation testing
2. A CCA-weight SM and connectome edge correlation of approximately *r=0.87*
3. One prominent CCA mode, defined as the primary mode’s (i.e. the strongest mode) z-scores for connectomes and SMs were at least a factor of 2 and 3 greater, respectively, than the next largest z-scores for connectomes and SMs (from any of the other modes)

The definition of a successful replication in the HCP 1200 dataset was:

1. At least one CCA mode that explains a statistically significant amount of variance in the connectomes and subject measures, relative to the null distribution generated via permutation testing
2. A positive and statistically significant CCA-weight SM and connectome edge correlation
3. One prominent CCA mode, defined as the primary mode’s (i.e. the strongest mode) z-scores for connectomes and SMs were at least a factor of 2 and 3 greater, respectively, than the next largest z-scores for connectomes and SMs (from any modes)

Criterion #2 of the HCP 1200 replication was selected to account for the larger amount of variability likely present in the HCP 1200 dataset. The criteria for success focus on the major findings of the original study, rather than attempting to replicate all numerical values exactly.

## 2. Methods

The scripts to run these analyses are available on the project’s Open Science Foundation page (https://osf.io/qm49a/).

### 2.1. Part 1 - HCP 500 Replication

#### 2.1.1. Data Download

The HCP 500 Parcellation, Timeseries, and Netmats dataset (hereafter referred to as HCP500), used in the original study, were obtained from *Connectome DB* (https://db.humanconnectome.org/app/action/ChooseDownloadResources?project=HCP_Resources&resource=GroupAvg&filePath=HCP500_Parcellation_Timeseries_Netmats.zip).

We used local, archived copies of the SMs (behavioral and demographic, also referred to as the “behavioral” and “restricted” files in our code, respectively), which were obtained from the HCP 500 subject release (https://www.humanconnectome.org/study/hcp-young-adult/document/500-subjects-data-release) when they first became available in 2014. These specific original files are no longer distributed on Connectome DB), however, the behavioral and phenotypic SM data for the HCP 1200 subject release are available and encapsulates the subjects in the HCP500 replication. We have not tested for differences in the data for the 461 subjects in our study between the releases, but others wanting to replicate this study can utilize the HCP1200 SM datasets (pulling the relevant data for the 461 subjects) if they do not have archived copies of the SM files from the original HCP500 release. Note that obtaining the restricted file requires additional permissions (https://www.humanconnectome.org/study/hcp-young-adult/document/wu-minn-hcp-consortium-restricted-data-use-terms).

#### 2.1.2. Data Preparation

Since the code provided by the original authors (https://www.fmrib.ox.ac.uk/datasets/HCP-CCA/) is solely the CCA analysis script, it does not prepare the functional connectivity and SM data for the analysis. Based on the instructions provided by the original authors to construct data matrices suitable for input into their Matlab CCA analysis script, we wrote a data preparation pipeline using Python and shell scripting (outlined in Figure 1).

**FIGURE 1:**
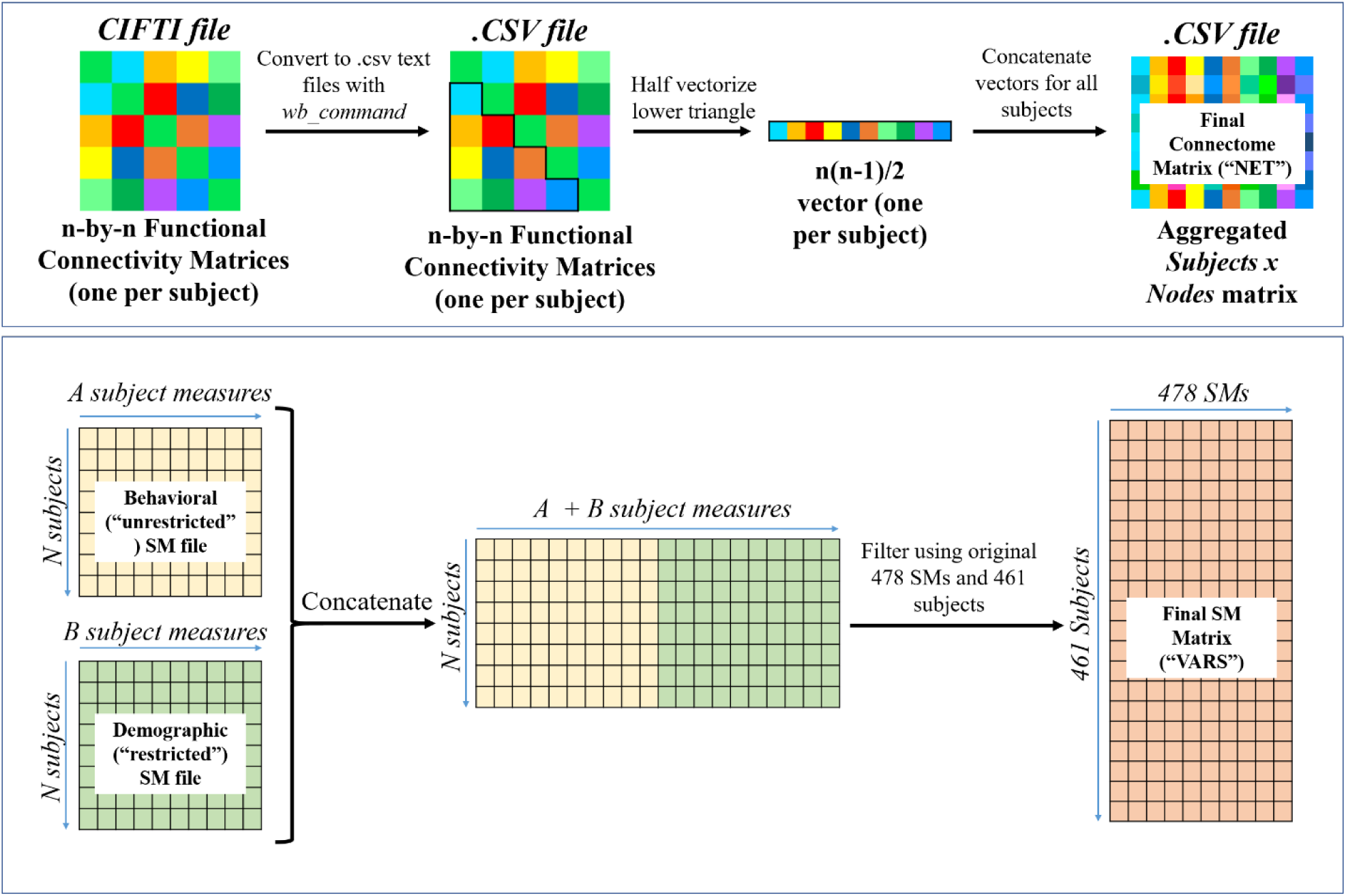
Preparation of the subject connectomes and SMs. **(Top)** In the general case, each subjects’ connectome (in CIFTI format) is first converted to comma-delimited text files (.csv); half-vectorization of the lower triangle is then performed on each subjects’ connectome, and the vectors for each subject are concatenated to form the final connectome matrix (“NET”). **(Bottom)** The behavioral and phenotypic SM files are concatenated, and then filtered based on the 461 subjects and 478 SMs used in the original study to generate the final SM matrix called (“VARS”).

Note that a preliminary analysis of the SM data revealed that multiple subjects were missing family relationship data pertinent to later permutation calculations. Consequently, since permutations would not be able to be generated using the *hcp2blocks* package, these subjects would have been dropped, and thus fewer than 461 subjects would have been included in our CCA analysis. To correct this, and ensure that we used all 461 subjects as in the original study, we wrote a ‘patch’ script that pulled the missing family relationship data for these subjects from the SM data in the HCP 1200 release.

Prior to constructing the SM and connectome matrices for the CCA, there was some pre-processing of the raw behavioral and phenotypic SM data files in order to factor binary and categorical variables, outlined in Table 1. The exact SM variable name is provided in parentheses.

**TABLE 1:** Numerical factoring of qualitative variables applied by Smith et al. 2015.

| *SM* | *Value (number or string)* | *Replacement* |
| --- | --- | --- |
|  | Blank cells | NaN |
|  | True/False | 1/0 |
| Reconstruction Software Version<br>(*fMRI_3T_ReconVrs*) | r227 | 1 |
|  | Blank/anything other than “r227” | 0 |
| Release (*Release*) | Q<1,2,3,...> | <1,2,3...> (remove the ‘Q’, keep the number) |
|  | MEG2 | 8 |
|  | S500 | 6 |
|  | S900 | 12 |
|  | S1200 | 16 |
| Gender (*Gender*) | Female/Male | 1/0 |
| Race (*Race*) | White | 0 |
|  | Black or African Am. | 1 |
|  | Asian/Nat. Hawaiian/Othr Pacific Is. | 2 |
|  | Am. Indian/Alaskan Nat. | 3 |
|  | More than one | 4 |
|  | Unknown or Not Reported | NaN |
| Ethnicity (*Ethnicity*) | Not Hispanic/Latino | 0 |
|  | Hispanic/Latino | 1 |
|  | Unknown or Not Reported | NaN |
| Vision (*Color_Vision*) | NORMAL | 0 |
|  | DEUTAN | 1 |
|  | TRITAN | 2 |
|  | TRITIAN | 3 |
|  | PROTAN | 4 |

The subjects and SMs that were included in our analysis are identical to those listed by the original authors. To arrive at the final list of SMs, the following quantitative inclusion criteria were applied to the SM data. A SM was kept only if *all* the criteria were met:

1. There was enough data available
  a. Defined as at least 50% of subjects having data for a given SM
2. There was sufficient variation in the SM
  a. Defined as less than 95% of subjects having the same SM value
3. The SM did not contain extreme outlier values based on the most extreme value from the median
  a. Specifically, a subject measure contained extreme outlier values if: *max(Ys) > 100*mean(Ys)*, where *Xs* is a vector of all subjects’ values for an SM *s*, and vector *Ys = (Xs - median(Xs))*^*2*^

Our replication did not require any pre-processing of resting-state fMRI data, since the resting-state scan data in the HCP 500 release is pre-processed (noise removal, motion correction, removal of structural artefacts, group-ICA derivation, and subject-level functional connectivity calculation) (S. M. Smith et al., 2015a). The 200-dimension functional connectivity matrices (i.e. the connectome data) used in the analysis are included in the HCP500 release as *CIFTI* files (https://www.nitrc.org/projects/cifti/). For a complete overview of the resting-state fMRI preprocessing steps and how the functional connectivity data were derived, please refer to the original study (S. M. Smith et al., 2015a). Because this data was supplied in CIFTI format, the Connectome Workbench (https://www.humanconnectome.org/software/connectome-workbench*) wb_command* tool was used to convert the CIFTI files to comma-delimited connectivity matrices.

Based on a list of 461 subjects provided by the original authors (see https://www.fmrib.ox.ac.uk/datasets/HCP-CCA/), the subject-level functional connectivity matrices were combined to create a large *Subjects x Edges* matrix. This was accomplished by vectorizing only the lower triangular part of each subject’s symmetric functional connectivity matrix (200 dimensions) to obtain a vector of length *n(n-1)/2* (equal to 19,900 for n=200), and concatenating these vectors to produce the final *Subjects x Edges* matrix (called “NET”) with dimensions 461×19900 (the general case is depicted in Figure 1, top). Using the list of SMs (478 total SMs) and subjects provided by the original authors, the SMs for each subject were similarly concatenated into a large *Subjects x SMs* matrix (named “VARS”) with dimensions 461×478 (Figure 1, bottom). The NET and VARS matrices are saved in comma delimited text files, named NET.txt and VARS.txt, respectively.

Preparation of the NET and VARS matrices can be performed using the *NET.py* and *VARS.py* scripts in our repository.

#### 2.1.3. Data Analysis

The entire CCA analysis pipeline is based on the code provided by the original authors at their website (https://www.fmrib.ox.ac.uk/datasets/HCP-CCA/) and the steps described in the original study [Smith 2015]. We outline the analysis steps here for clarity. Prior to the actual CCA itself, there are several pre-processing steps (outlined in the flowchart, Figure 2).

**FIGURE 2:**
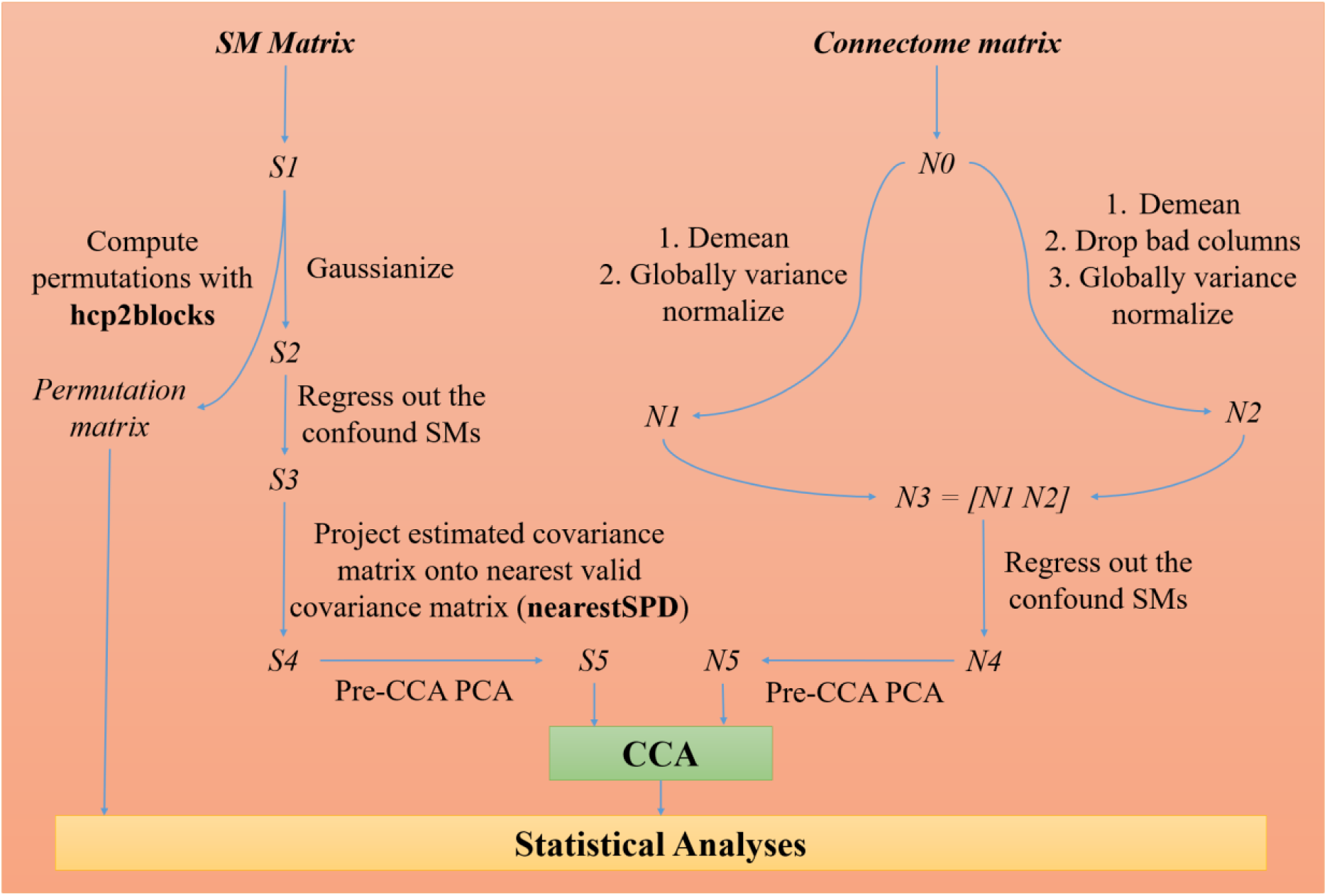
A general outline of the data analysis stage, highlighting the pre-processing stages for subject measures (**LEFT SIDE**) and connectomes (**RIGHT SIDE**) immediately prior to the CCA.

The SM data was imported into Matlab from the VARS.txt file generated in the Data Preparation stage; 10,000 permutations (preserving family structure) were generated using the package *hcp2blocks* (https://github.com/andersonwinkler/HCP/blob/master/share/hcp2blocks.m). Note that in the original study 100,000 permutations were used, however we could only perform 10,000 permutations at the time of conducting this replication due to computational limitations. An updated version of this pre-print with 100,000 permutations will be uploaded in the future.

The subject measure matrix was initially stored in matrix *S*_*1*_. *S*_*2*_ was formed by applying quantitative inclusion criteria (programmatically) to matrix *S*_*1*_, dropping SMs that do not meet all inclusion criteria, and then applying gaussian normalization to the matrix. After application of the inclusion criteria, 158 SMs remained in matrix *S*_*2*_.

Matrix *S*_*3*_ was then generated by regressing out the 17 confound variables identified in the original study. There were 9 confound variables identified in the original study (the actual SM variable name is given in parentheses):

1. Acquisition reconstruction software version (*quarter/release)*)
2. Average subject mead motion (*rfMRI_motion)*
3. Weight (*weight*)
4. Height (*height*)
5. Systolic blood pressure (*BPSystolic*)
6. Diastolic blood pressure (*BPDiastolic*)
7. Hemoglobin A1C (*HbA1C*)
8. Cube-root of total brain volume (*FS_BrainSeg_Vol*)
9. Cube-root of total intracranial volume (*FS_InterCranial_Vol*)

In addition to these 9, additional confounds were created by demeaning and squaring SMs 2-9 in order to account for any potential nonlinear effects of those confounds, for a total of 17 confounds. All confounds were demeaned, and any missing data were imputed as zero.

Matrix *S*_*4*_ was created by projecting an estimated covariance matrix onto the nearest valid covariance matrix, using the Matlab package nearestSPD. Finally, *S*_*4*_ was input to a 100-dimension PCA, generating matrix *S*_*5*_, which was the SM data matrix input to the CCA.

The connectomes were processed in a similar manner - first, the connectome data was imported into Matlab from the NET.txt file generated in the Data Preprocessing stage, forming matrix *N*_0_. From *N*_0_, two matrices (*N*_1_ and *N*_2_) were generated, and then horizontally concatenated to form matrix *N*_3_. *N*_1_ was formed by first demeaning *N*_0_, then variance normalizing it. *N*_2_ was formed by demeaning *N*_0_, followed by dropping any badly formatted columns (columns whose mean values were very low, z-score<0.1), and finally variance normalizing it. The 17 confound variables identified in the original study were then regressed out of *N*_3_, forming matrix *N*_4_. *N*_4_ served as the input to a 100-dimension PCA to reduce dimensionality, estimating the top 100 eigenvectors of the connectomes, forming matrix *N*_5_, which was the connectome data matrix input to the CCA.

The CCA was run using the Matlab *canoncorr* function (from the Statistics and Machine Learning Toolbox), by using the 70-eigenvector matrices for the SMs (*S*_*5*_) and connectomes (*N*_*5*_) as inputs. The CCA estimated 100 modes, optimizing de-mixing matrices A and B such that the resulting matrices *U=N5*A* and *V=S*_*5*_*\*B* (both had dimensions 7,810×70) were maximally similar to each other. By correlating the corresponding column pairs in *U* and *V* matrices (i.e. the same mode in each), we determined the strength for which a mode of covariation exists in both the subject connectomes and subject measures. By regressing matrix *U* against *V*, we determined an overall Pearson’s *r* value for the correlation between the CCA modes.

To estimate the significance of the correlation between *U* and *V*, we performed a 10,000 permutation-based significance test in which null distributions were calculated using the subject permutations to calculate thresholds for significance in the percent variance explained in the SMs and connectomes by each CCA mode. The 5th and 95th percentiles of the null distribution were used to determine which of the first 20 CCA modes explained a significant percentage of the total variance in the original SM (*S*_*1*_) and connectome (*N*_*0*_) data matrices. Connectomes and subject measures were analyzed separately.

### 2.2. Part 2 - HCP 1200 Replication

#### 2.2.1. Data Download

The HCP 1200 dataset, used in the original study, was obtained from the *Connectome DB* (https://db.humanconnectome.org/app/action/ChooseDownloadResources?project=HCP_Resources&resource=GroupAvg&filePath=HCP1200_Parcellation_Timeseries_Netmats.zip). Note that to obtain the HCP dataset and associated subject measures, users need to request access permission on the *Connectome DB* website (db.humanconnectome.org).

We also obtained copies of the subject measures (behavioral and demographic, referred to as the “behavioral” and “restricted” files in our code, respectively) for the HCP 1200 subject release, from the Connectome DB.

#### 2.2.2. Data Preparation

Although a head motion summary measure is not included in either the behavioral or phenotypic SM data files for the HCP 1200 release, these values can be obtained in the full HCP 1200 dataset (https://db.humanconnectome.org/app/action/ChooseDownloadResources?project=HCP_Resources&resource=GroupAvg&filePath=HCP_S1200_PTNmaps_d200.zip). A head motion summary was obtained by averaging the four *MovementRelativeRMSmean* measurements for all four of a subject’s resting-state scans (available in the *Movement_RelativeRMS_mean.txt* values for the rfMRI runs, found in each subject’s */MNINonLinear/Results/{rfMRI_REST1_LR, rfMRI_REST1_RL, rfMRI_REST2_LR, rfMRI_REST2_RL}* folders).

The preparation of subject measures to form the *Subject x Edges* and *Subjects x SMs* matrices was performed analogously to the process outlined in section *2.1.4*.

The HCP 1200 release includes a comma-delimited text file (called *netmats2.txt*) whose rows are the entire symmetric functional connectivity matrix (from the 200-dimension group ICA, with functional connectivity calculated using FSLNets partial correlation with Tikhonov regularization, rho=0.01) for each individual subject, vectorized (i.e. 200×200 matrix flattened into a 40,000 element vector). The derivation of these connectomes matched the process applied in the original study (S. M. Smith et al., 2015a).

The supplied *netmats2.txt* file had dimensions 1003×40,000; after performing the half-vectorization for each subject (after initially reconstructing their 200×200 matrices) the NET matrix had dimensions 1003×19900.

#### 2.2.3. Data Analysis

The analysis of the HCP1200 data was performed identically to that of the HCP500 data. After applying the quantitative exclusion criteria to the SMs, 149 remained in the analysis to be fed into the PCA to generate matrix S5.

## 3. Results

The results of the replications are summarized in Table 2.

**TABLE 2:** Results of the computational replications using the HCP 500 and HCP 1200 datasets.

|  | *Original Study* | *HCP500 Replication* | *HCP1200 Replication* |
| --- | --- | --- | --- |
| *Permutations* | 100,000 | 10,000 | 10,000 |
| *Correlation of Mode 1 weights* | $r=0.87$<br>$p<10^{-5}$ | $r=0.8715$<br>$p<10^{-4}$ | $r=0.7432$<br>$p<10^{-4}$ |
| *Percentage Variance Explained (Z-score)* |  |  |  |
| *Connectome* | 0.53% (7.7) | 0.526% (7.428) | 0.366% (5.796) |
| *SMs* | 1.7% (9.2) | 1.749% (9.307) | 1.963% (10.708) |
| *Next highest Z-scores* |  |  |  |
| *Connectome* | 2.7 | 2.8101 | 3.424 |
| *SMs* | 2.4 | 2.7275 | 6.672 |

The results of the HCP 500 replication are shown in Figure 3.

**FIGURE 3:**
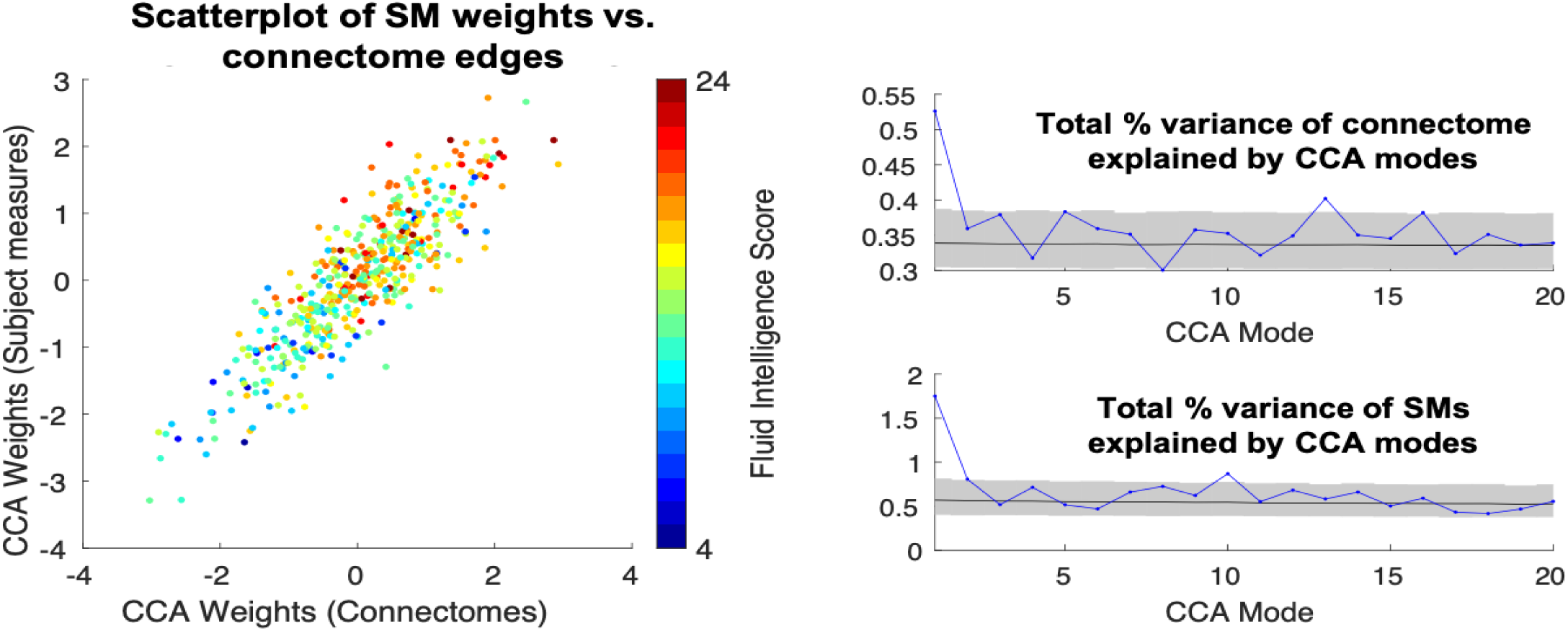
(**left**)Scatter plot of the SM weights against connectome weights from mode 1 of the CCA using HCP 500 data, with each one point per subject, and subject’s fluid intelligence score indicated by colors. The total percent of variance in connectomes and subject measures as explained by the first 20 CCA modes. (**right**) CCA mode 1 explained the largest portion of variance in both. Shaded portions indicate the 5th and 95th percentiles of the null distribution, with the mean of the distribution shown as a black line.

The results of the HCP1200 replication are shown in Figure 4:

**FIGURE 4:**
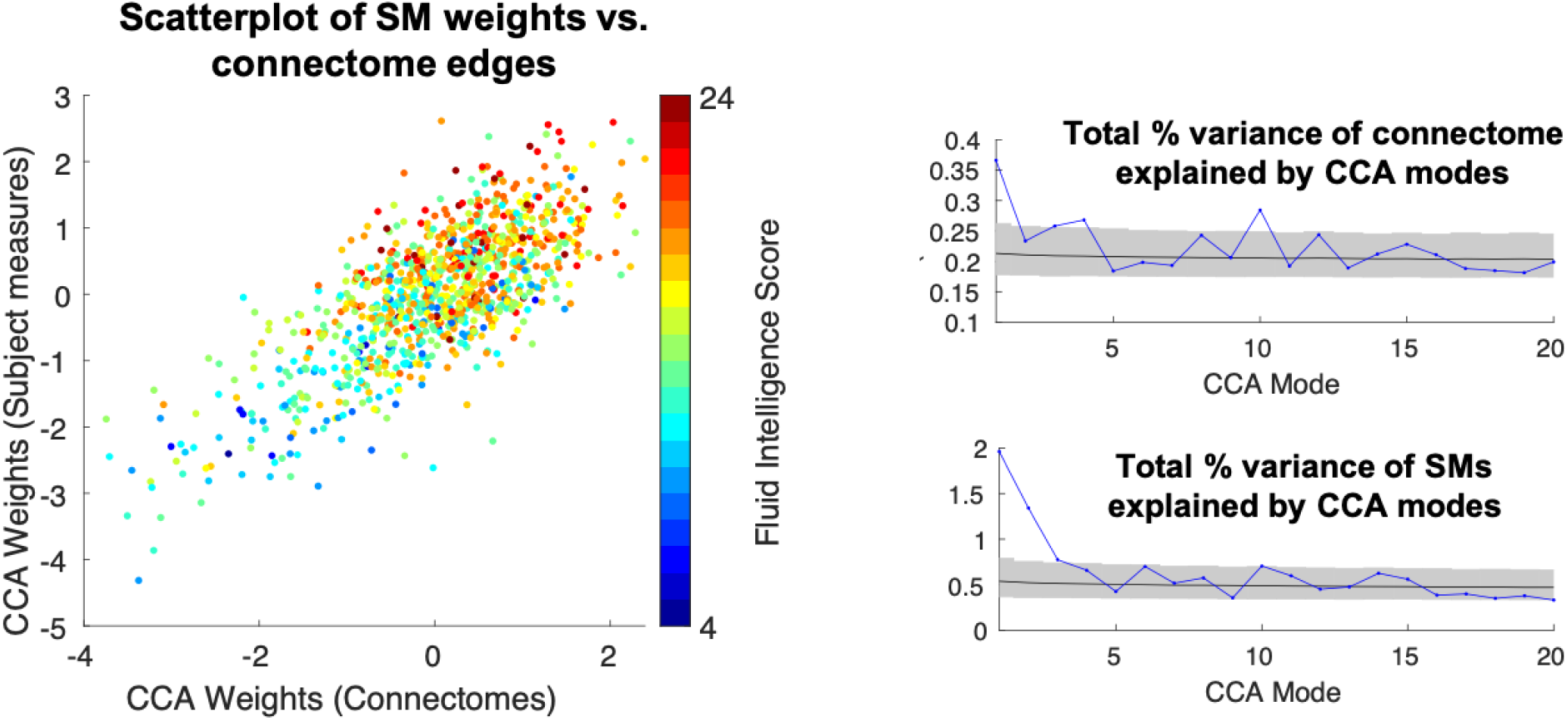
(**left**) Scatter plot of the SM weights against connectome weights from mode 1 of the CCA using HCP 1200 data, with each one point per subject, and subject’s fluid intelligence score indicated by colors. (**right**) CCA mode 1 explained the largest portion of the total percentage of variance in both connectomes and subject measures. Shaded portions indicate the 5th and 95th percentiles of the null distribution, with the mean of the distribution shown as a black line.

In both replications, only 10,000 permutations were possible due to computational limitations, whereas in the original study 100,000 permutations were used. As a result, the significance of the correlations is p<10^−4^.

## 4. Discussion

### 4.1. HCP 500 Replication

The computational replication in the HCP 500 dataset was successful according to the criteria stated in the introduction.

- Criteria 1 was fully met:

- The primary mode explained a significant amount of variance in the connectomes (0.526%) and SMs (1.749%). These percentages were nearly identical to those reported in the original study of 0.53% for connectomes and 1.7% for SMs.
- Criteria 2 was fully met:

- There was a correlation of *r=0.8715* (p<10^−4^) between the primary CCA mode connectome and SM weights. Note that the significance level was p<10^−4^ whereas in the original study it was p<10^−5^. This was due only 10,000 permutations being calculated.
- Criteria 3 was fully met:

- The primary CCA mode z-score for connectomes was 7.428 (near the original author’s value of 7.7) and was a factor of 2 larger than the next highest z-score of 2.8101.
- The primary CCA mode z-score for SMs was 9.307 (also near the originally reported value of 9.2) and was a factor of 3 larger than the next highest z-score of 2.7275.

***Given that the results of the exact computational replication are nearly identical to the major findings reported by Smith et al***., ***we conclude that our analysis pipeline properly replicates the original***. The minor differences in the results are likely due to a) small differences in our copy of the SM data, and b) only conducting 10,000 permutations, whereas the original study used 100,000. An updated version of this pre-print with 100,000 permutations will be uploaded in the future.

### 4.2. HCP 1200 Replication

The computational replication in the HCP 1200 dataset was partially successful according to the criteria established in the introduction.

- Criteria 1 was met:

- The primary mode explained a significant amount of variance in the connectomes (0.366%) and SMs (1.963%) relative to the null distribution.
- Criteria 2 was met:

- There was a correlation of r=*0.7432* (p<10^−4^) between the primary CCA mode connectome and SM weights. Note that the significance level was p<10^−4^ whereas in the original study it was p<10^−5^. This was due only 10,000 permutations being calculated.
- Criteria 3 was only partially met:

- The primary CCA mode z-score for connectomes was 5.796 and ***was not*** a factor of 2 larger than the next highest z-score of 3.424.
- The primary CCA mode z-score for SMs was 10.708 and ***was not*** a factor of 3 larger than the next highest z-score of 6.672.

Since this analysis was an extension of the original, it has no implications on the validity of our pipeline. However, the results of this analysis do support our hypothesis that the original study results would appear in the HCP 1200 dataset given its similarity to the HCP 500 dataset. Interestingly, with the larger dataset we found that the primary CCA mode explained a *smaller* portion of the variance in connectomes (0.366% in HCP 1200, vs. ∼0.53% in HCP 500), and a *larger* percentage of the variance in SMs (1.963% in HCP 1200, vs. ∼1.7% in HCP 500) when compared to the original analysis in the HCP 500 dataset.

### 4.3. Conclusion

In conclusion, this computational replication was successful in not only replicating the results of the original study, but also in validating our data preparation and CCA analysis pipeline for use in future extensions of the original study utilizing independent non-HCP datasets.

## Acknowledgements

The authors are supported by the NIMH Intramural Program ZICMH002960.

## HCP Data

HCP500 and HCP1200 data were provided [in part] by the Human Connectome Project, WU-Minn Consortium (Principal Investigators: David Van Essen and Kamil Ugurbil; 1U54MH091657) funded by the 16 NIH Institutes and Centers that support the NIH Blueprint for Neuroscience Research; and by the McDonnell Center for Systems Neuroscience at Washington University.

## HPC

This work utilized the computational resources of the NIH HPC Biowulf cluster (http://hpc.nih.gov).

